# Isolation of phosphate solubilizing *Pseudomonas* strains from apple rhizosphere in the Trans Himalayan region of Himachal Pradesh, India

**DOI:** 10.1101/193672

**Authors:** Ranjna Sharma, Joginder Pal, Mohinder Kaur

**Author notes:** Indicates the corresponding author (Ranjna Sharma).

## Abstract

Total fifteen phosphate solubilizing bacteria were isolated from rhizosphere soil of apple tree on Kings’ B medium belonging to genus *Pseudomonas* spp. They were characterized on the basis of morphological and biochemical characteristics. Preliminary selection as phosphate solubilizing bacteria were done on the basis of formation of transparent zone around the colony on Pikovskaya’s agar medium containing 0.5% triclcium phosphate (TCP). Maximum *in vitro* phosphate solubilization on Pikovskaya’s agar plates after 72h incubation at 28°C was shown by An-15-Mg (46 mm), whereas in case of broth, again this strain showed maximum tricalcium phosphate (TCP) solubilization (76 μg/ml). The pH of each inoculated broth was recorded daily and dropped significantly (pH 7.0-3.99). All the *Pseudomonas* isolates were further evaluated for overall plant growth promoting traits. Greatest siderophore activity was exhibited by An-14-Mg (71.23 %SU) followed by An-15-Mg (70.12 %SU) which were statiscally at par with each other. Whereas, maximum IAA production was observed again in An-15-Mg (95 μg/ml). This isolate also showed the maximum production of HCN and ammonia. 16S rDNA and phylogenetic analysis showed that strain An-15-Mg exhibits 99% level of similarity with *Pseudomonas aeruginosa.* Therefore, it was designated as *Pseudomonas aeruginosa* strain An-15-Mg. HPLC analysis showed that the *Pseudomonas aeruginosa* strain An-15-Mg produced maximum concentration of succinic acid, malonic, citric and malic acid with small amounts of schimic, quinic, tartaric, fumaric and lactic acids. The strain An-15-Mg possessed phosphate solubilization as major PGP trait along with different PGP traits. The potential phosphate solubilizing strain of *Pseudomonas aeruginosa* was reported first time as PGPR which lives in close association with apple tree without harming the plant, therefore could be used as a promising phosphate solubilizer and biofertilizer in apple crop grown in high hills of Himachal Pradesh.

**Authors contribution:** All authors made significant efforts towards the completion of this research work and preparation of manuscript timely. Author Ranjna Sharma has done collection of soil samples of apple tree from two districts, isolation and characterization of phosphate solubilizing *Pseudomonas* isolates and their screening for different PGPT’s, phenotypic characteriazation, 16S rRNA analysis and statistical analysis. Author S P Singh has done HPLC analysis at their institute. Author Ankita has done sequence analysis and phylogenetic tree preparation. Author Mohinder Kaur is my mentor and generated idea regarding this work. She guided me in completing this work successfully as well as manuscript writing.

## Introduction

Phosphorus is an essential macronutrient required for the growth of the plants (Illmer and Schinner 1992). The phosphorus is applied to soil in the form of phosphatic fertilizers. 75–90% of phosphatic fertilizer when added to soil is precipitated by metal cation complexes (Stevenson 1986). It has been suggested that the accumulated phosphates in agricultural soils are sufficient to sustain maximum crop yields worldwide for about 100 years (Goldstein et al. 1993). The excessive application of P causes environmental and health problems. Therefore, it is necessary to have alternate solution to overcome this problem. Phosphate-solubilizing abilities of soil microorganisms have proved to be an economically sound alternative to the more expensive phosphatic fertilizers.

The organisms with phosphate-solubilizing potential increase the availability of soluble phosphate and can enhance plant growth by increasing the efficiency of biological nitrogen fixation or enhancing the availability of other trace elements such as iron, zinc, etc., and by production of plant growth-promoting regulators (Sattar and Gaur 1987; Kucey et al. 1989; Ponmurugan and Gopi 2006).

Solubilization and mineralization of phosphorus compounds by PGPR in soil seems to be one of the possible pathways by which plants takes inorganic phosphorus for their growth and development. 98% of Indian soils are deficient in phosphorus, because the concentration of free phosphorus, the form available to plants even in fertile soils is generally not higher than 10 μM even at pH 6.5 where it is most soluble (Narsian and Patel 2009).

Although microbial inoculants are in use for improving soil fertility during the last century, however, a meager work has been reported on P solubilization compared to nitrogen fixation. Microbial biomass assimilates soluble P, and prevents it from adsorption or fixation (Khan and Joergensen 2009). Microorganisms enhance the P availability to plants by mineralizing organic P in soil and by solubilizing precipitated phosphates (Pradhan and Sukla 2005; Chen et al. 2006). These bacteria in the presence of labile carbon serve as a sink for P by rapidly immobilizing it even in low P soils (Bünemann et al. 2004).

Several machanisms may be involved for solubilization of insoluble phosphorus but, the major one is the action of organic acids synthesized by soil microorganisms (Mishra et al. 1990). Production of organic acids results in acidification of the microbial cell and its surroundings. Consequently, inorganic phosphorus (Pi) may be released from a mineral phosphate by proton substitution for Ca^+2^ (Goldstein 1994).

Fluorescent *Pseudomonas* sp. are effective phosphate solubilizers which directly or indirectly promote the growth of plants by increasing the concentration of available nutrients and antibiosis. Therefore, the aim of the present study was to explore the potential phosphate solubilizing PGP *Pseudomonas* sp. associated with apple tree in temperate and sub temperate zones of Himachal Pradesh. All the strains were tested for i*n vitro* production of phosphate solubilization and other PGP activities. The potential phosphate solubilizing *Pseudomonas* strain has been used for organic acid estimation through HPLC and its molecular analysis was done.

## Material and Methods

### Collection of Soil Samples

Soil samples were collected at 5–15cm depth in the area of tree basin. A total of Twelve composite samples of soil were drawn from normal (Seven) and diseased sites (five) of apple from Shimla and Kullu Districts of Himachal Pradesh, India and used for isolation of phosphate solubilizing bacterial (PSB) population. Samples were immediately stored at 4°C in plastic bags loosely tied to ensure sufficient aeration and to prevent moisture loss until isolation of bacterial community. Isolates from normal sites were designated as An whereas isolates from sick site as Ar.

### Isolation of phosphate solubilizing rhizobacteria

Phosphate solubilizing rhizobacteria were isolated from collected soil samples of apple trees by serial dilution using a standard spread plate technique on Pikovskaya’s (PVK) medium (Pikovskaya 1948) supplemented with tri-calcium phosphate (TCP, 0.5%) as an insoluble inorganic phosphate source. PSB were expressed as colony forming unit (CFU) per gram of dry soil weight. These isolated colonies were maintained on PVK medium with 30% glycerol on cryopreservation tubes at -20°C for further study.

### Phenotypic Identification

Further identification of Phosphate solubilizing rhizobacterial population isolated from collected soil samples of apple trees were done on the basis of Gram staining, spore staining, cellular morphology and other biochemical tests as per their genera as prescribed in Bergey’s Manual of Systematic Bacteriology. The most predominant rhizobacterial isolates showing greenish/yellowish fluorescent pigments at 302 nm wavelength in BIO-RAD gel doc system XR were assumed to be *Pseudomonas* sp.

### Phosphate Solubilization

All the isolates were grown in PVK broth (10 ml) supplemented with 0.5% tri-calcium phosphate (TCP) for 72 hours at 28±2°C under shake conditions (100 rpm). Supernatant was prepared by centrifugation of culture at 12,000 rpm for 20 minutes and stored at 4°C. 100 μl cell free supernatant of each isolate was applied to 10 mm wells of pre poured sterilized PVK agar plates (Pikovsakaya 1948). Plates were incubated at 28±2°C for three days and observed for P solubilisation in the form of clear zone produced around the well. Further quantitative estimation of phosphorus was done in PVK broth by the vanadomolybdate method (Sundara Rao and Sinha 1963). Change in pH of the culture broth was recorded by pH metre equipped with a glass electrode.

### Screening of phosphate solubilizing bacteria for other plant growth promoting traits

#### Siderophore activity

Siderophore production was observed in terms of reduction in blue color as per cent siderophore units (% SU) by the method (Schwyn and Neilands 1987).

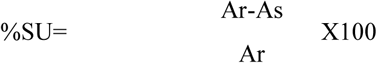

Ar=Absorbance of reference at 630 nm, As=Absorbance of supernatant at 630 nm

### IAA production

Quantitative estimation of IAA was carried out by the method described by Gordon and Paleg (1957).

### HCN and Ammonia production

Hydrogen cyanide (HCN) production was determined on King’s B agar medium supplemented with 4.0g/l glycine after 3 days of incubation at 28±2°C (Bakker and Schippers 1987). Whereas ammonia production was determined by the method given by Lata and Saxena 2003.

### Identification of potential phosphate solubilizing *Pseudomonas* strain and its phylogenetic analysis

The rhizobacterial isolate that was identified as the potential phosphate solubilizing *Pseudomonas* isolate during *in vitro* evaluation was characterized to species level assessed for organic acid production by HPLC. Bacterial genomic DNA was isolated by using DNA purification kit (Banglore Genei Pvt. Ltd., India) as per the manufacturer’s instructions. *Pseudomonas* specific primer set (forward 5’ GGTCTGAGAGGATGATCAGT 3’ and reverse 5’ TTAGCTCCACCTCGCGGC3’) was used to amplify 16S rDNA of potential phosphate solubilizer. PCR reactions were carried out in 20ml reaction volume containing, 50 ng of template DNA, 20pmoles of each primer, 0.2mM dNTPs and 1U Taq polymerase in 1 × PCR buffer. Reactions were performed under following conditions: initial denaturation at 94° C for 30s, 588C for 1 min. followed by 35 cycles of 92° C for 1 min., annealing at 55° C for 1min 30s and final extension at 72°C for 10min. The amplified PCR product was separated by gel electrophoresis on 1.0% (w/v) agarose gel. The purified fragment was sequenced from Xcleris lab, Ahmadabad, India using gene specific primers (both forward and reverse primers). The sequence obtained were subjected to database search using BLAST nsuite (NCBI). Based on 16S rRNA gene sequences, phylogenetically related bacteria were aligned using a BLAST search (Altschul et al. 1997). Multiple alignment with sequences of related taxa of the genus *Pseudomonas* was implemented using CLUSTAL W (version 1.6). The neighbour-joining phylogenetic tree was constructed and analyzed using MEGA 6 software. The topology of the phylogenetic tree was evaluated by the bootstrap resampling method of Felsenstein (1985) with 100 replicates. The sequence was submitted to NCBI GenBank database and accession number was assigned.

For the analysis of organic acids, *Pseudomonas* strain An-15-Mg was grown and multiplied in PVK liquid medium. A 2 ml of three day old cell free supernatant of *Pseudomonas* strain An-15-Mg obtained after solubilization of tri-calcium phosphate (0.5%) was homogenized using a vortex mixer and then centrifuged at 10,000 × g at 4°C for 10 min. The supernatant was filtered through 0.2 μm syringe filter and 20 μl filtrate was injected into HPLC-MS/MS system. The samples in autosampler were kept at 4°C.The organic acids were separated by using a Hi Plex H (7.7 × 300 mm × 8 μm), (Agilent Technologies Pvt. Ltd., Chandigarh, India) column and a guard column (Hi–Plex H 3 × 5mm×8 μm) (Agilent Technologies) maintained at 60°C at a flow rate of 0.6 mL/min. The samples were run isocratically using 0.1 % formic acid in water for 20 min. The MS/MS analysis was performed with a hybrid triple quadrupole/ion trap mass spectrometer, QTRAP 5500 (ABSciex India Pvt. Ltd., Gurgaon, India). The mass spectra were acquired using TurboIonSpray ionization in negative ion mode and scheduled MRM using analyst software.

The compound–dependent MS parameters were determined by infusion of each compound. The curtain gas was adjusted to 30 psi. The ion spray voltage, ion source gas 1, and ion source gas 2 were – 4.5 kV, 50 psi, and 50 psi, respectively. The temperature of the source was fixed to 550°C. The authentic standards of organic acids were used for preparation of calibration curve. The organic acids in the samples were determined by comparing the retention times and peak areas of chromatograms with the standards for malic acid, malonic acid, citric acid, tartaric acid, succinic acid, formic acid, lactic acid, quinic acid and schimic acid. The quantification of organic acids was conducted using MultiQuant software.

### Statistical Analysis

Experimental data were analysed using standard analysis of variance (ANOVA) followed by Duncan’s multiple comparison tests (p < 0.05). Standard errors were calculated for all mean values. Arcsine transformation was applied to data expressed in percentage.

## Results

### Morphological and biochemical characteristics of phosphate solubilizing bacterial isolates

The phosphate solubilizing bacterial isolates were isolated from collected soil samples of both the districts on Pikovskaya’s agar medium. The bacterial colonies were circular/irregular in shape, raised elevation, entire edge with transparent opacity producing bluish/greenish to brown pigment on nutrient agar plates. They were identified biochemically as positive for catalase, oxidase and oxidative metabolism, whereas negative for Grams staining, tween 80 hydrolysis and spore staining. 80% of the isolates were positive for denitrification test, whereas 66.67% isolates were positive for Gelatin liquefaction test. Fifteen bacterial isolates showed optimum growth at 28°C on King’s B medium (Table 1). On the basis of physiological and biochemical characteristics, the P-solubilizing bacteria were identified as *Pseudomonas* spp.

**Table:1.** Physiological and biochemical characteristics of bacterial isolate from rhizosphere soil of normal and sick sites of apple

| District | Site | Isolate | Characteristics |  |  |  |  |  |  | Growth<br>(King's B<br>agar)<br>28°C |
| --- | --- | --- | --- | --- | --- | --- | --- | --- | --- | --- |
| Shimla | Normal |  | Gram Staining | Tween 80<br>Hydrolysis | Metabolism<br>(oxidative) | Denitrifi-Cation | Oxidase | Caalase | Gelatin<br>Liquifaction |  |
|  |  | An-2-Mg | - | - | + | - | + | + | - | + |
|  |  | An-3-Mg | - | - | + | + | + | + | + | + |
|  |  | An-4-Mg | - | - | + | + | + | + | + | + |
|  |  | An-5-Mg | - | - | + | - | + | + | - | + |
|  |  | An-6-Mg | - | - | + | - | + | + | - | + |
|  |  | An-7-Mg | - | - | + | + | + | + | + | + |
|  |  | An-9-Mg | - | - | + | + | + | + | + | + |
|  |  | An-10-Mg | - | - | + | + | + | + | + | + |
|  | Sick | Ar-1-Mg | - | - | + | + | + | + | + | + |
|  |  | Ar-8-Mg | - | - | + | + | + | + | - | + |
|  |  | Ar-11-Mg | - | - | + | + | + | + | - | + |
| Kullu | Normal | An-12-Mg | - | - | + | + | + | + | + | + |
|  |  | An-13-Mg | - | - | + | + | + | + | + | + |
|  |  | An-14-Mg | - | - | + | + | + | + | + | + |
|  |  | An-15-Mg | - | - | + | + | + | + | + | + |

(+) indicates positivity and (-) negativity of test

### Screening of phosphate solubilizing bacteria for plant growth promoting traits (PGPTs)

All the fifteen *Pseudomonas* isolates were screened for plant growth promoting traits (Table 2). The *Pseudomonas* isolate An-15-Mg showed highest P-solubilization both in plate assay (46 mm) and liquid assay method (79 μg/ml) simultaneousely maximum decrease in the pH of the medium (pH 7.0-3.99). In vitro evaluation of biocontrol features showed that highest per cent siderophore unit (% SU) was shown by the isolate An-14-Mg (71.23 %SU) followed by An-15-Mg (70.12 %SU) which were significantly at par with each other after 72 h of incubation at 28°C. Significantly higher auxin production was exhibited by two *Pseudomonas* isolate An-7-Mg and An-15-Mg (95 μg/ml each). Greatest HCN production was exhibited by isolate An-15-Mg (++++) followed by An-7-Mg (+++), whereas maximum ammonia production was obtained in four *Pseudomonas* isolates Ar-1-Mg, Ar-11-Mg, An-14-Mg and An-15-Mg (++++).

**Table 2.**
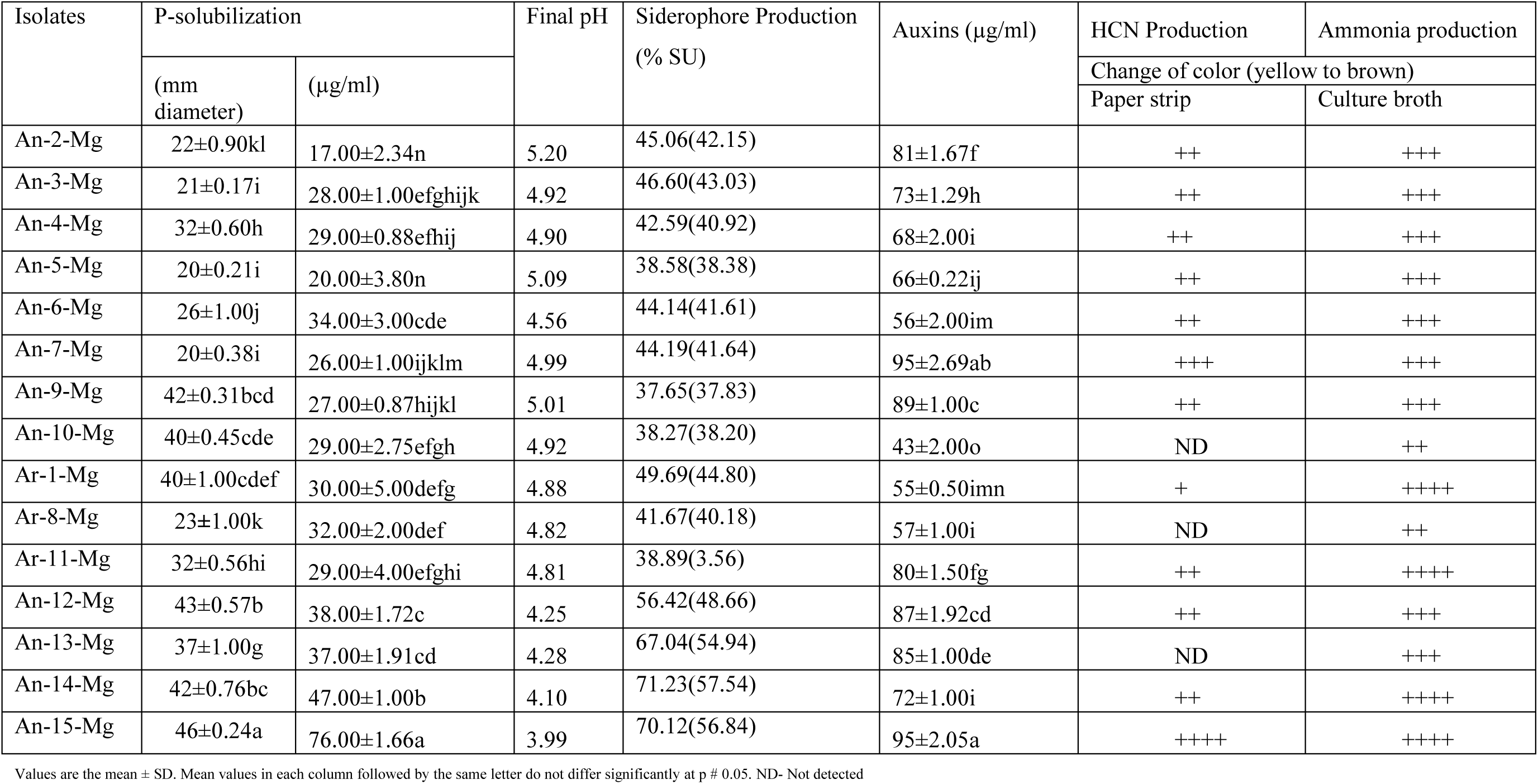
Different plant growth promoting traits of P-solubilizing *Pseudomonas* isolates

### Phylogenetic analysis of potential *Pseudomonas* strain An-15-Mg by 16S rDNA sequence analysis

The *Pseudomonas* strain An-15-Mg was found to be the most effective phosphate solubilzer under *in vitro* conditions along with other plant growth promoting traits. Therefore the strain An-15-Mg was identified to the species level by 16S rDNA sequence analysis. The 16S rDNA fragment was successfully amplified using *Pseudomonas* specific primers and sequenced. Both the forward and reverse sequences so obtained were corrected, and unreadable and ambiguous sequences were deleted. Finally a complete sequence of 787bp was obtained. Sequence analysis revealed that isolate An-15-Mg showed 99% homology with *Pseudomonas aeruginosa* (CP017353). A phylogenetic tree also verified the identity of strain An-15-Mg as *P. aeruginosa* because the isolate was closely clustered with *P. aeruginosa* (CP015002) high boot strap value of 65% (Figure 2). The sequence of strain An-15-Mg was deposited in NCBI Genbank under accession number KJ500026 and was designated as *Pseudomonas aeruginosa* strain An-15-Mg.

### Production of different organic acids

HPLC-MS/MS system analysis of culture filtrate showed the presence of multiple organic acids during the solubilization of tricalcium phosphate. A decrease in the pH of the growth medium was observed after 72h (pH 7.0-pH 3.99). Among the different authentic organic acids used for the comparison, production of succinic acid (1.65μg/ml) was higher followed by malonic acid (0.390μg/ml), citric acid (0.378 μg/ml) and malic acid (0.162 μg/ml) given in Table 3 and Figure 2. Quinic acid (0.089μg/ml), lactic acid (0.086 μg/ml), tartaric acid (0.065 μg/ml) and schimic acid (0.020 μg/ml) and fumaric acid (0.013 μg/ml) were produced in small amount during the solubilization of tricalcium phosphate.

**Table 3.**
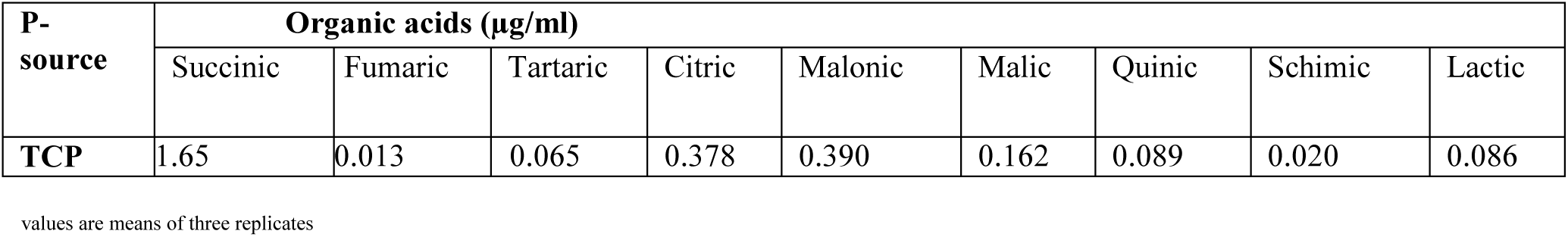
Organic acid production during solubilization of phosphate substrate (TCP) by *Pseudomonas aeruginosa* strain An-15-Mg after three days of incubation at 28±2°C

## Discussion

In the present study, total fifteen phosphate solubilizing plant growth promoting bacteria belonging to genus *Pseudomonas* spp. were isolated from healthy and sick sites of apple tree on King’s B medium from Shimla and Kullu districts of Himachal Pradesh. The preliminary selection as phosphate solubilizer were done on Pikovskaya’s agar medium. The yellow coloured colonies surrounded by transparent halo were selected and purified on the same medium. They all belongs to *Pseudomonas* spp. isolates on the basis of morphological and biochemical characterization (Table 1). *Pseudomonas* spp. a well-known plant-associated bacteria and effective root colonizer detected in this study as abundant members of the cultivable P-solubilizing PGPR community. This is most likely due to the microbial community composition of the study sites.

Phosphate solubilization is the important mechanism of plant growth promotion. Insufficient availability of phosphorus limits the growth of plants. The low solubility of tricalcium phosphate, hydroxyapatite and aluminum phosphate causes low availability of phosphorus in apple orchards. All the *Pseudomonas* isolates were screened for phosphate solubilization and other plant growth promoting traits (PGPTs). They all produce atleast one or more plant growth promoting traits. Maximum production of phosphate solubilization under plate and liquid assay method was shown by An-15-Mg (Table 2).

Park et al. 2009 showed that *Pseudomonas fluorescens* RAF15 produced a wide range of secondary metabolites like siderophore, HCN, and IAA which showed potential efficacy not only against phytopathogenic fungi but also promote plant growth. The role of siderophore in biocontrol of several fungal phytopathogens has been reported (Scher and Baker 1982). Several plant growth-promoting rhizobacteria are known to secrete IAA into culture media, and have been shown to stimulate plant growth (Gaudin et al. 1994). In addition to insoluble P solubilizaion by producing different organic acids substantial production of siderophore, HCN, ammonia and IAA by potential phosphate solubilizing strain *Pseudomonas aeruginosa* An-15-Mg clearly suggests its use as biofertilizer and phosphate solubiizer.

Several mechanisms have been proposed for the suppression of phytopathogens by antagonistic bacteria, including production of antibiotics, siderophore and lytic enzymes such as amylase, chitinase, protease and cellulase (Amaresan et al. 2012).

Multiple biocontrol mechanisms have been implicated in the suppression of fungal root diseases by biocontrol species of Bacillus. It has been strongly suggested that the main success of a biocontrol agent is largely attributable to multifunctional biocontrol traits (Vassilev et al. 2006).

16S rRNA gene sequence analysis is an authentic technique used to study bacterial isolate at species level and allows their identification as well as prediction of phylogenetic relationship (Naz and Bano 2010). Sequence analysis of potential phosphate solubilizing *Pseudomonas* isolate An-15-Mg revealed that it showed 99% similarity with *Pseudomonas aeruginosa* (CP017353). A phylogenetic tree also verified the identity of strain An-15-Mg as *P. aeruginosa* because the isolate was closely clustered with *P. aeruginosa* (CP017353) high boot strap value 65% (Figure 1).

**Figure 1.**
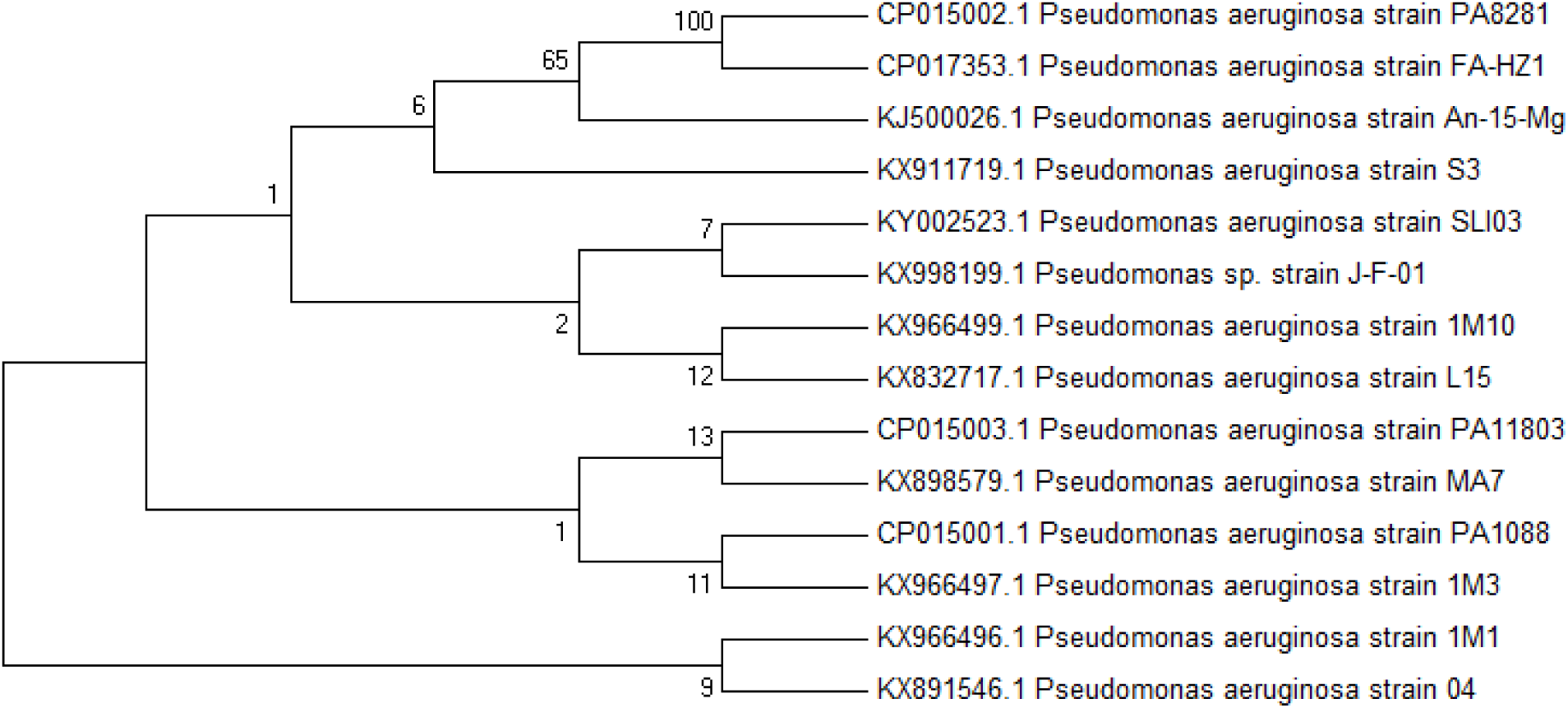
Neighbour-joining phylogenetic dendogram based on 16S rRNA sequences showing relationships between isolate An-15-Mg and related taxa.

Exudation of nutrients from roots of apple tree might exert a selective pressure on the proliferation of plant growth promoting microbial community such as *Pseudomonas* and *Bacillus*. Selective pressure through different host plants has been suggested earlier by Siciliano and Germida 1999, as observed in the present study.

HPLC analysis of 72h old culture supernatant of *Pseudomonas aeruginosa* mainly produced succinic, malonic, citric, malic, oxalic acids with small amount of lactic, schimic, tartaric and quinic acids in 72h old culture supernatant containing glucose as major substrate (Figure 2), Simultaneousely there were observed a decrease in the pH of the medium from pH 7.0-3.99. Goldstein et al. 1993, showed that organic acids like glycolic, gluconic, succinic, oxalic, citirc and malonic acids have been identified in phosphate solubizers namely *Bacillus firmus, Pseudomonas cepacia and Pseudomonas* sp. A decline in the pH of the medium during solubilization of phosphate substrates suggested the secretion of organic acids by *Acinetobacter rhizosphaerae* BIHB 723 as reported for other bacteria (Chen et al. 2006). Gulati et al. 2010, showed that the strain *Acinetobacter rhizosphaerae* BIHB 723 produced different organic acids during the solubilization of phosphate substrates explicating the influence of substrate on the production of organic acids i.e oxalic, gluconic, 2-Keto gluconic, lactic, formic and malic acid. The higher solubilization of TCP being due to its amorphous nature and is more facile to solubilization.

**Figure 2.**
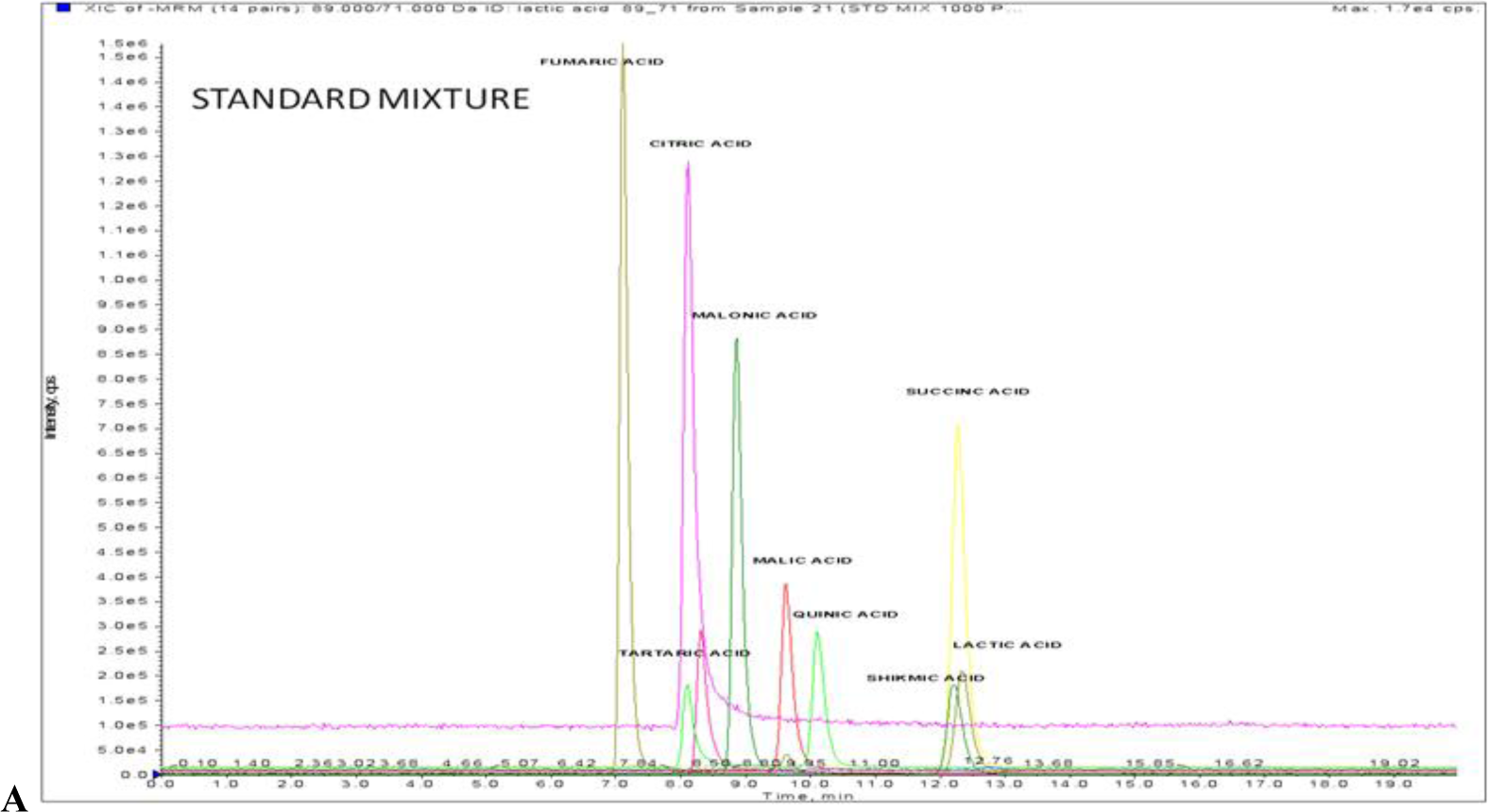
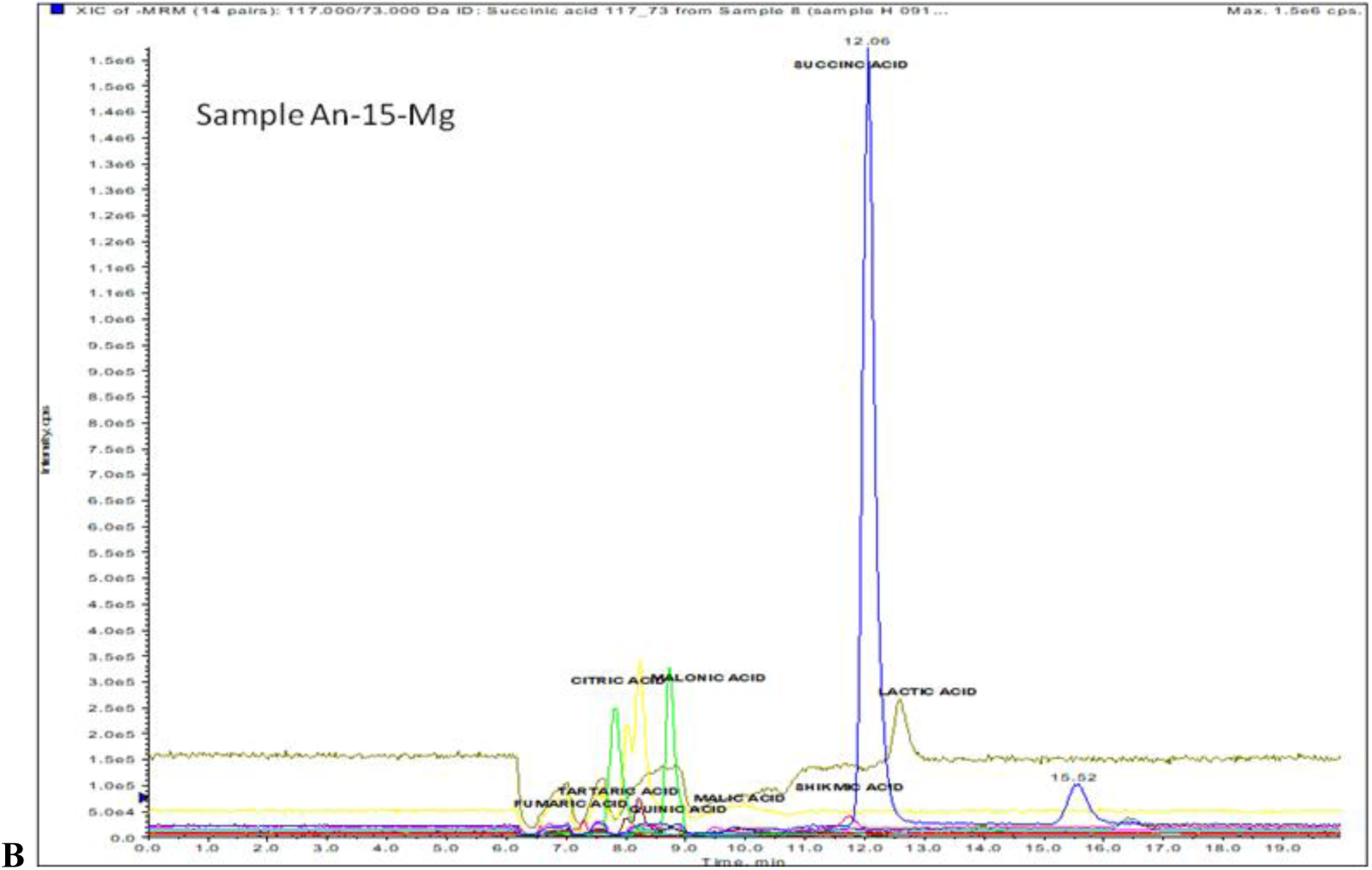
HPLC chromatograms of authentic standards (A) and different organic acids produced by potential P-solubilizing *Pseudomonas aeruginosa* strain An-15-Mg (B)

## Conclusion

Fluorescent *Pseudomonas* sp. are commonly found in rhizosphere of different crops. Potential phosphate solubilizing *Pseudomonas aeruginosa* strain An-15-Mg isolated from the rhizospheric soil of apple tree could be used as a potential phosphate solubilizer in agricultural environments. Because of the innate potential of producing siderophore, IAA, HCN and ammonia, *Pseudomonas aeruginosa* strain An-15-Mg could be used as a biofertilizer as well as a potential biocontrol agent. *In vitro* production of different PGPTs cannot always be reproduced under field conditions. Therefore, field evaluation of this strain as effective plant growth promoting bacterium is still under process.

## Acknowledgement

The present study was supported by funds from the Department of Science and Technology, New Delhi and Department of Basic Sciences, Dr. YSP UHF, Nauni, Solan Himachal Pradesh, India.

